# Stem cell factor and granulocyte colony-stimulating factor promote remyelination in the chronic phase of severe traumatic brain injury

**DOI:** 10.1101/2023.01.24.525450

**Authors:** Xuecheng Qiu, Suning Ping, Michele Kyle, Lawrence Chin, Li-Ru Zhao

**Affiliations:** Department of Neurosurgery, State University of New York Upstate Medical University, Syracuse, New York, 13210, USA

**Keywords:** Traumatic brain injury, chronic phase, SCF, G-CSF, demyelination, remyelination

## Abstract

Severe traumatic brain injury (TBI) causes long-term disability and death in young adults. White matter is vulnerable to TBI damage. Demyelination is a major pathological change of white matter injury after TBI. Demyelination which is characterized by myelin sheath disruption and oligodendrocyte cell death leads to long-term neurological function deficits. Stem cell factor (SCF) and granulocyte colony–stimulating factor (G-CSF) treatments have shown neuroprotective and neurorestorative effects in the subacute and chronic phases of experimental TBI. Our previous study has revealed that combined SCF and G-CSF treatment (SCF+G-CSF) enhances myelin repair in the chronic phase of TBI. However, the long-term effect and mechanism of SCF+G-CSF-enhanced myelin repair remain unclear. In this study, we uncovered persistent and progressive myelin loss in the chronic phase of severe TBI. SCF+G-CSF treatment in the chronic phase of severe TBI enhanced remyelination in the ipsilateral external capsule and striatum. The SCF+G-CSF-enhanced myelin repair is positively correlated with the proliferation of oligodendrocyte progenitor cells in the subventricular zone. These findings reveal the therapeutic potential of SCF+G-CSF in myelin repair in the chronic phase of severe TBI and shed light on the mechanism underlying SCF+G-CSF-enhanced remyelination in chronic TBI.

## 1. Introduction

Traumatic brain injury (TBI) causes a primary traumatic axon injury and secondary injury in white matter tracts [1]. TBI has become a major cause of death and long-term disability among children and young adults [2]. Epidemiological data on TBI from the Centers for Disease Control and Prevention of the United States reveal about 2.87 million TBI-related emergency department visits and hospitalizations in 2014 including 56,800 TBI-related deaths [3]. Approximately 10–15% of patients with TBI have more severe injuries and require long-term care [4], which causes a high economic burden.

Growing clinical evidence has shown long-term white matter impairments in TBI patients [5–8], even in mild TBI patients [9]. Myelin is a major component of white matter. The myelin sheath enwrapping axons is comprised of oligodendrocyte processes. Post-TBI white matter injury involves both traumatic axon injury (TAI) and myelin pathology. Myelin is vulnerable to damage from TAI after TBI [8]. Persistent accumulation of myelin debris after TAI has been found in a rat model of TBI [10]. In TBI patients, chronic inflammation in the white matter with white matter degeneration persists years after TBI [11]. Disrupted myelin-released myelin-associated proteins such as Nogo, oligodendrocyte myelin glycoprotein, and myelin-associated glycoprotein negatively mediate axonal regeneration and neural plasticity after injury [12]. Numerous studies have shown that TBI induces widespread oligodendrocyte death and demyelination, followed by concomitant oligodendrogenesis and remyelination [13–16]. Manipulating oligodendrogenesis and remyelination may provide a novel therapeutic target to promote white matter integrity and improve neurological function recovery after brain injury [17,13].

Stem cell factor (SCF) and granulocyte-colony stimulating factor (G-CSF) were originally discovered as hematopoietic growth factors to regulate and maintain hematopoiesis [18,19]. Both SCF and G-CSF can pass through the blood-brain barrier in intact animals [20,21], revealing the possibility of SCF and G-CSF in regulating physiological function in the brain. Increasing evidence has shown that SCF and G-CSF can enhance angiogenesis, neurogenesis, neurite outgrowth, neuronal network remodeling, and neuronal function in neurological disorders [22–24]. It has been reported that SCF and G-CSF treatments attenuate white matter loss and increase the proliferation of intrinsic oligodendrocyte precursor cells in animal models of spinal cord injury [25,26]. Our previous study has also revealed that the combined treatment of SCF and G-CSF (SCF+G-CSF) in the chronic phase of severe TBI enhances remyelination and improves neurological functional recovery [27]. However, the time-dependent effects of SCF+G-CSF treatment in promoting myelin repair in the chronic phase of severe TBI remains unclear.

In the present study, young adult mice with severe TBI received 7-day treatments of SCF+G-CSF initiated at 3 months post-TBI. The myelin changes in the ipsilateral white matter were assessed 1 day, 21 weeks, and 36 weeks after SCF+G-CSF treatment. The possible mechanism of the SCF+G-CSF-enhanced myelin repair in the chronic phase of severe TBI was also explored.

## 2. Materials and Methods

All procedures of animal experiments were approved by the Institutional Animal Care and Use Committee of SUNY Upstate Medical University and performed in accordance with the National Institutes of Health Guide for the Care and Use of Laboratory Animals.

### 2.1. Controlled Cortical Impact Model of TBI

The controlled cortical impact (CCI) TBI model was used in this study for making a severe TBI model in young adult mice as described in our previous studies [23] [27]. Briefly, C57BL/6J male mice (The Jackson Laboratory, USA) at the age of 12 weeks were anesthetized with Avertin (Sigma-Aldrich, 0.4 g/kg body weight, i.p.). The heads of mice were fixed on a stereotaxic frame, and a 4-mm-diameter cranial window centered at 2 mm lateral to the bregma on the right side of the skull was created. The brain was impacted using a 3-mm diameter flat tip with a 4° medial angle, 2-mm depth strike from the surface of the dura, at a 1.5 m/sec strike velocity and 8.5 second contact time. As sham controls, mice were handled in the same manner but without craniotomy and impact. After surgery, mice were placed on a homeothermic blanket (37°C) to prevent post-anesthesia hypothermia. An antibiotic (ampicillin, 20-100 mg/kg, s.c.) and analgesic (sustained-release buprenorphine, 0.6 mg/kg, s.c.) were injected before and after surgery respectively.

### 2.2. Experimental design

To carry out a time course experiment for assessing the therapeutic effects of SCF+G-CSF on myelin repair in the chronic phase of TBI, a total of 43 male mice were divided into three-time points: 1 day, 21 weeks, and 36 weeks after SCF+G-CSF treatment. Each time point has three groups: a sham group, a TBI-vehicle control group, and a TBI-SCF+G-CSF treatment group. The SCF+G-CSF treatment was initiated 3 months after TBI, corresponding to the chronic phase of TBI. Mice were subcutaneously injected daily with recombinant mouse SCF (200 μg/kg/day, diluted with saline, PeproTech) and recombinant human G-CSF (50 μg/kg/day, diluted with 5% dextrose, Amgen), or an equal volume of vehicle solution for 7 days according to our previous TBI study [23]. The experimental flowchart is shown in Figure 1. Mice were euthanized at the selected time points: 1 day (sham, n = 5; TBI-vehicle, n = 4; TBI-SCF+G-CSF, n =5), 21 weeks (sham, n = 6; TBI-vehicle, n = 6; TBI-SCF+G-CSF, n =6) and 36 weeks (sham, n = 3; TBI-vehicle, n = 4; TBI-SCF+G-CSF, n =5) post-treatment. Mice with health problems during the experiment were excluded from this study.

**Figure 1.**
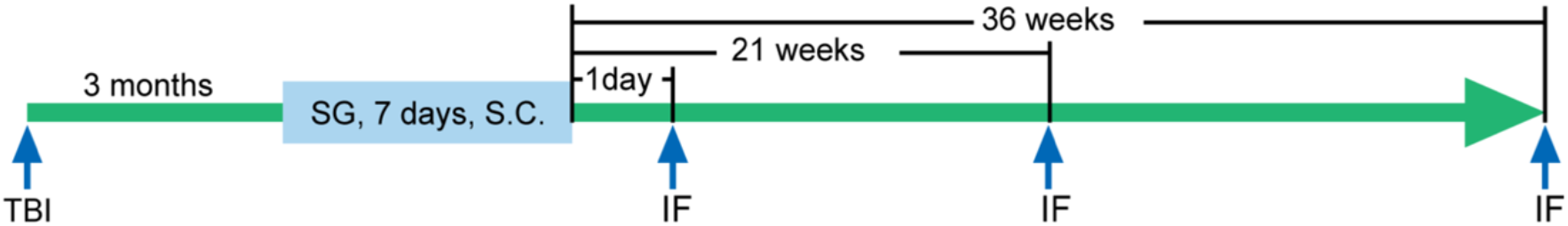
A schematic flowchart of the experiment. The mice were treated with SCF+G-CSF (SG) for 7 days at 3 months after TBI. The immunofluorescence (IF) staining on brain sections was performed at 1 day, 21 weeks, and 36 weeks post-treatment, respectively.

### 2.3. Immunofluorescence staining

Mice were anesthetized with Ketamine (90mg/kg) and Xylazine (9mg/kg) (i.p.) and transcardially perfused with phosphate-buffered saline (PBS) containing heparin (10U/ml, Sagent Pharmaceuticals) followed by 10% neutral buffered formalin (Sigma) perfusion. Brains were removed, post-fixed in the same fixative solution overnight at 4°C and placed in 30% sucrose solution in PBS. After being fully dehydrated, the brains were embedded in O.C.T. compound (Thermo Fisher Scientific) on dry ice and sectioned at a thickness of 30 μm using a Leica cryostat.

For free-floating immunofluorescence staining, brain sections were washed with PBS (pH 7.4) and incubated in PBS with 10% normal donkey serum, 0.3 % Triton X-100 (Sigma-Aldrich), and 1% IgG-free bovine serum albumin (BSA) (Sigma-Aldrich) for 1 hour at room temperature to block non-specific binding. Brain sections were incubated with the primary antibodies including rabbit anti-MBP (1:500, Abcam), rabbit anti-Ki67 (1:500, ThermoFisher), goat anti-Olig2 (1:500, Novus Biologicals), and mouse anti-Sox2 (1:500, ThermoFisher) overnight at 4°C. After rinsing with PBS, brain sections were incubated with the corresponding secondary antibodies including donkey anti-rabbit IgG conjugated with Alexa fluor 488 or 594 (1:300, Invitrogen), donkey anti-goat IgG conjugated with Alexa fluor 488 or 594 (1:300, Invitrogen) or donkey anti-mouse IgG conjugated with Alexa fluor 488 or 594 (1:300, Invitrogen) for 2 hours at room temperature in the dark. After rinsing with PBS, brain sections were mounted on Superfrost Plus Slides (Thermo Fisher Scientific) with VECTASHIELD HardSet Antifade Mounting Medium (Vector Laboratories). Immunofluorescence-positive staining was captured with a confocal microscope (Zeiss LSM780) through a 40x objective. The number of Olig2^+^ or Olig2^+^/Sox2^+^ cells was counted using ImageJ software (NIH software). The percentage of the MBP^+^ area was also analyzed using ImageJ software.

### 2.4. Culture of oligodendrocyte progenitor cells isolated from adult mouse brain

Adult C57BL/6J mice (male, 3-4 months old) were decapitated after being anesthetized with Ketamine (90mg/kg) and Xylazine (9mg/kg) (i.p.). The brains were removed quickly and placed into ice-cold Hibernate E (Thermo Fisher Scientific). The forebrain was dissected, and the meninges were mechanically removed. The forebrain was then mechanically minced into 1 mm^3^ pieces. The tissue pieces were spun down by a centrifuge at 100 g for 1min. The pellet was mixed with 10 mL of dissociation solution (0.2% papain containing 1.1 mM EDTA, 0.067 mM mercaptoethanol and 5.5 mM cysteine-HCl in the DMEM high glucose medium) and incubated for 60 min at 37 °C on a shaker (50 rpm). The digestion was stopped by adding ice-cold Hanks’ Balanced Salt Solution (HBSS). The mixture was centrifuged at 200 g for 3 min at room temperature. The pellet was resuspended in 10 mL ice-cold Hibernate E medium. To obtain a single cell suspension, the tissue suspension was triturated 10 times using a 5-mL serological pipette and subsequently three 1-mL pipette tips 10 times. The tissue suspension was then allowed to sediment for 1 min, and the supernatant was filtered through a 70 μm cell strainer to obtain the single-cell suspension. To get more cells, 5 mL of fresh ice-cold Hibernate E medium was added to the sediment, and the above procedures were repeated. The collected cell suspension was mixed with isotonic Percoll (GE Healthcare, 90% Percoll 10% 10x PBS, pH 7.2) at a final concentration of 22.5% and centrifuged at 800 g for 20 min without break at room temperature. The myelin debris and supernatant were discarded. The brain cell-containing phase (about the last 2 mL) and cell pellet were resuspended in PBS, transferred to a fresh 15 mL tube, and centrifuged at 300 g for 5 min at room temperature. The supernatant was discarded. The cell pellet was resuspended in red blood cell lysis buffer (BD Bioscience) and incubated for 1 min at room temperature to remove red blood cells. Thereafter, 10 mL of modified washing buffer (MWB) containing 2 mM EDTA, 2 mM Na-Pyruvate, and 0.5% BSA in D-PBS (pH 7.2) was added to the cell suspension and spun down using a centrifuge at 300 g for 5 min at room temperature. The cell pellets were resuspended in 0.1 mL of MWB. To acquire purified OPCs, anti-A2B5 microbeads (Miltenyi Biotec) were used for magnetic-activated cell sorting according to the manufacturer’s instructions. The sorted A2B5 positive OPCs were flushed out of the column with pre-warmed, CO2 and O2 pre-equilibrated OPC growth medium containing DMEM high glucose medium (Thermo Fisher Scientific), 60 μg/mL N-Acetyl cysteine (Sigma-Aldrich), 10 μg/mL human recombinant insulin (Thermo Fisher Scientific), 1mM sodium pyruvate (Thermo Fisher Scientific), 50 μg/mL apo-transferrin (Sigma-Aldrich), 16.1 μg/mL putrescine (Sigma-Aldrich), 40 ng/mL sodium selenite (Sigma-Aldrich), 60 ng/mL progesterone (Sigma-Aldrich), 330 μg/mL bovine serum albumin (Sigma-Aldrich) supplemented with epidermal growth factor (EGF) (10 ng/mL, Thermo Fisher Scientific), basic-fibroblast growth factor (b-FGF) (20 ng/mL, Thermo Fisher Scientific) and platelet-derived growth factor AA (PDGF-AA) (20 ng/mL, Sigma-Aldrich) [28]. The OPCs were seeded onto PDL-coated (100 μg/ml, Sigma-Aldrich) wells of a 12-well plate at the concentration of 0.5 x10^6^ OPCs per well. The cell medium was half exchanged every 3 days and passaged after confluent. To identify the purity of the cultured OPCs, the passaged cells were seeded on PDL-coated coverslips for 24 hours, and immunostained with rabbit anti-NG2 (1: 300, Millipore), goat anti-Olig2 (1: 500, Novus Biologicals) and rabbit anti-Sox2 (1: 300, Millipore) using the above-mentioned routine immunofluorescence staining protocol. The cells were then imaged under a confocal microscope (LSM 780) using a 20 x Objective lens.

### 2.5. Proliferation and differentiation assay of oligodendrocyte progenitor cells

For proliferation assays, OPCs (passage 1) were cultured in the OPC growth medium with/without SCF+G-CSF (20 ng/mL) for 24 hours. BrdU (Roche, 20 ng/mL) was then added in the medium 6 hours before fixation. For immunofluorescence staining, the cells were fixed with 10% neutral buffered formalin for 10 min at room temperature. After washing with PBS, the cells were treated with 2N HCl for 30 min at room temperature to hydrolyze DNA for BrdU staining. After rinsing with PBS, the nonspecific staining was blocked with 10% normal donkey serum in PBS with 1% BSA and 0.3% triton for 1 hour and then incubated with mouse anti-BrdU (1: 1000, Roche), rabbit anti-Ki67 (1: 600, ThermoFisher), and goat anti-Olig2 (1: 500, Novus Biologicals) overnight at 4 °C. The next day, the cells were rinsed with PBS 3 times and incubated with donkey anti-mouse IgG conjugated with Alexa fluor 488, donkey anti-rabbit IgG conjugated with Alexa fluor 594, or donkey anti-goat IgG conjugated with Alexa fluor 647 at room temperature for 2 hours. After rinsing with PBS, coverslips were mounted with DAPI mounting medium and imaged with a confocal microscope (LSM 780) using a 20 x Objective lens.

For differentiation assays, OPCs (passage 1) were cultured in the OPC growth medium without all growth factors. OPCs were treated with medium alone as a vehicle control, triiodothyronine (T3, 40 ng/mL, Sigma-Aldrich) as a positive control, or SCF+G-CSF (20 ng/mL) for 48 hours. Cells were fixed with 10% neutral formalin solution for 10 min and rinsed with PBS three times. The fixed cells were immunostained with rabbit anti-MBP (1:500, Abcam). The cells were imaged with an Olympus microscope using a 20 x Objective lens.

### 2.6. Statistical analysis

Data are presented as the mean ± standard error of the mean (SEM). All statistical analyses were performed in GraphPad Prism (GraphPad Software, Inc.). Before the analysis of parametric data, Shapiro-Wilk tests were performed to ensure the normality of the data. For multiple groups, data were analyzed using either One-way or Two-way ANOVA followed by Tukey’s *post hoc* test or Fisher LSD tests and presented as significant differences when the P values reached statistical significance. Unpaired Student’s t-tests were used for two group comparisons. Two-tailed t-tests were used throughout, and statistical significance was defined as p < 0.05. For correlation analyses, we used Pearson’s correlation coefficient test (linear regression).

## 3. Results

### 3.1. SCF+G-CSF treatment increases remyelination in the chronic phase of severe TBI

Demyelination, which has been observed in both human TBI patients and rodent TBI models, has become a therapeutic target for TBI [17,13,8,29]. White matter tracts are highly vulnerable to brain injury [30]. It has been reported that oligodendrocyte apoptosis remains up to at least 3 months in the ipsilateral external capsule which is the subcortical region directly next to the cortical injury site in a mouse model of TBI [14]. Our previous study has revealed that post-TBI demyelination occurs in the ipsilateral striatum and hippocampus, and that SCF+G-CSF treatment shows superior efficacy in enhancing remyelination than that of SCF and G-CSF alone treatment in the chronic phase of severe TBI [27]. In the present study, we determined the time-course changes of demyelination in the ipsilateral external capsule and striatum and explored the timing effects of SCF+G-CSF treatment on myelin repair in the chronic phase of severe TBI.

Myelin basic protein (MBP), as a myelin marker, was used for detecting TBI-caused demyelination and SCF+G-CSF-induced remyelination in the chronic phase of severe TBI. Compared with the sham group, the percentage of MBP^+^ area was significantly decreased in the ipsilateral external capsule and striatum in all TBI mice including the mice in TBI-vehicle and TBI-SCF+G-CSF groups at 1 day, 21 weeks, and 36 weeks post-treatment (i.e. 13, 34, and 49 weeks after TBI, respectively) (Figure 2) (Figure 3 AC and E). In addition, in all TBI mice, the percentage of MBP^+^ area in both the ipsilateral external capsule and striatum was also significantly decreased at 49 weeks post-TBI compared with 34 weeks after TBI (Figure 2). These findings indicate that severe TBI causes persistent and progressive demyelination in the white matter.

**Figure 2.**
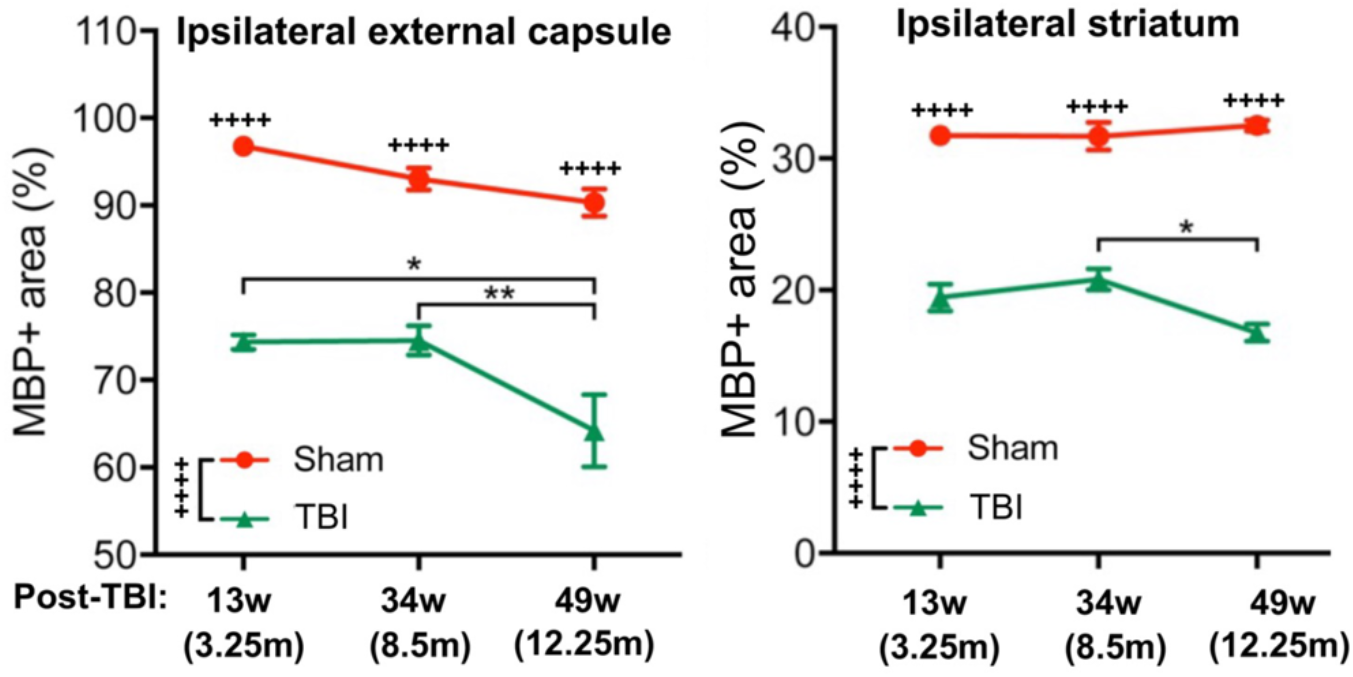
Persistent and progressive white matter demyelination in the ipsilateral external capsule and striatum 13, 34, and 49 weeks after TBI (i.e. at 1 day, 21 weeks, and 36 weeks post-treatment, or 3.25, 8.5, and 12.25 months after TBI). Myelin basic protein (MBP) immunostaining. Two-way ANOVA followed by Tukey *post hoc test*. Time point difference: *p<0.05, **p<0.01; Sham vs. TBI: ++++p<0.0001.

In the ipsilateral external capsule, the MBP^+^ area in the TBI-SCF+G-CSF group showed an increasing trend at 21 weeks post-treatment and significant increases at 36 weeks post-treatment compared to the TBI-vehicle group (Figure 3 A and C). In the ipsilateral striatum, the SCF+G-CSF treatment significantly increased the MBP^+^ area at 1 day and 36 weeks post-treatment and showed an increasing trend at 21 weeks posttreatment compared with the TBI-vehicle group (Figure 3 B and E). These findings indicate that SCF+G-CSF treatment enhances remyelination in the white matter during the chronic phase of severe TBI.

**Figure 3.**
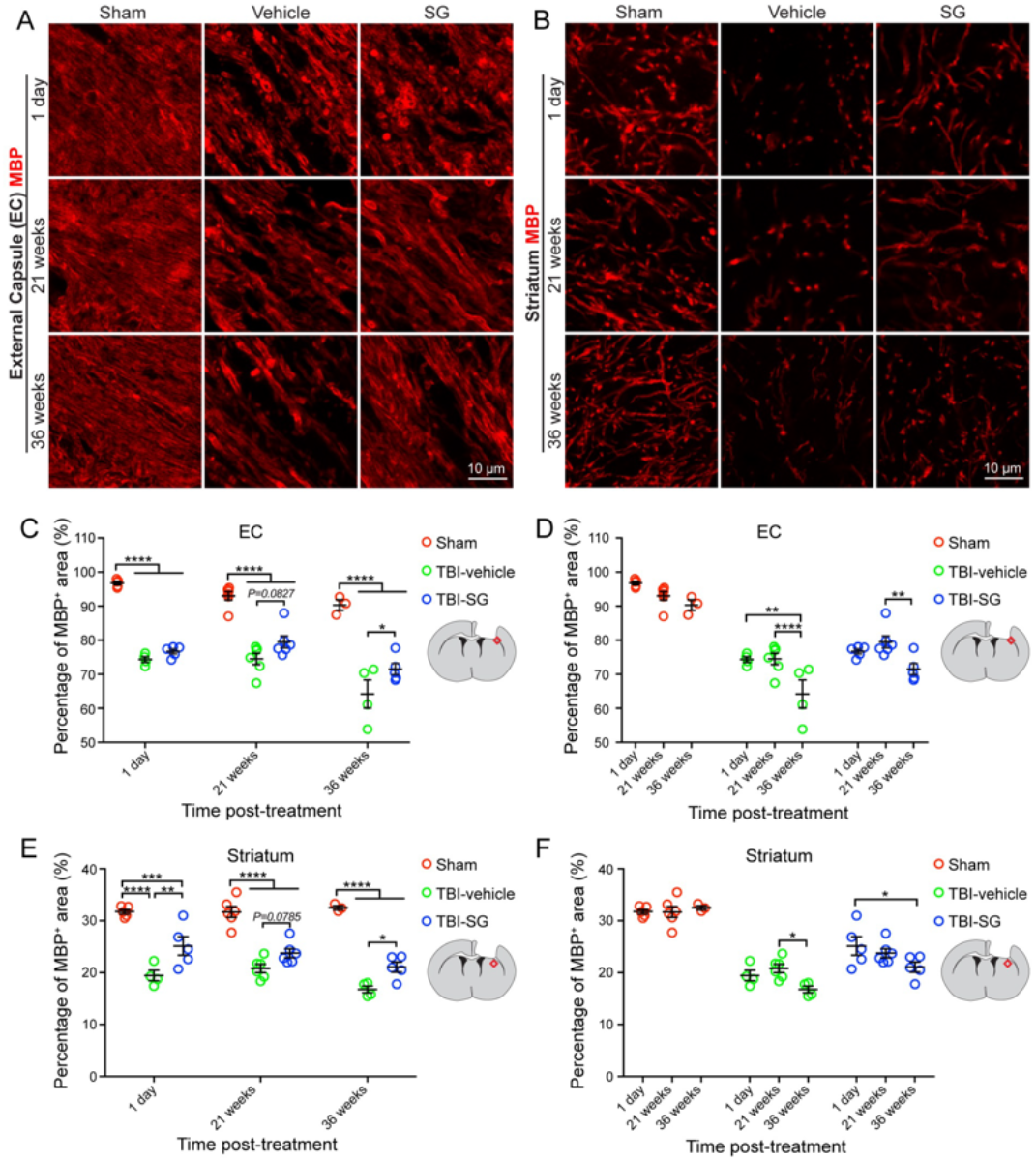
SCF+G-CSF treatment increases remyelination in the chronic phase of severe TBI. **(A** and **B)** Representative images of MBP immunopositive myelin in the ipsilateral external capsule (**A**) and striatum (**B**) at 1 day, 21 weeks, and 36 weeks post-treatment. **(C - F)** Quantitative data of analysis of the MBP immunopositive area in the ipsilateral external capsule **(C** and **D)** and striatum **(E** and **F**). Twoway ANOVA followed by Tukey *post hoc test*, \**p* < 0.05, \*\**p* < 0.01, \*\*\**p* < 0.001, \*\*\*\**p* < 0.0001. Scale bars: 10 μm. SG: SCF+G-CSF. EC: external capsule.

The time-course data showed that the MBP expression in both the ipsilateral external capsule and striatum had no significant differences between 1 day and 21 weeks after treatment (i.e. between 13 and 34 weeks after TBI) in both the TBI-vehicle and TBI-SCF+G-CSF groups (Figure 2) (Figure 3D and F), suggesting that progressive demyelination may not occur yet during this particular period after severe TBI. However, the MBP expression was significantly decreased at 36 weeks post-treatment (i.e. 49 weeks after TBI) in both the TBI-vehicle group and TBI-SCF+G-CSF group (Figure 2) (Figure 3D and F), indicating that the progressive demyelination happens in the late chronic phase of severe TBI. Altogether, severe TBI leads to long-term and progressive demyelination in the white matter. SCF+G-CSF treatment in the chronic phase of severe TBI shows a long-term effect in enhancing remyelination in the ipsilateral external capsule and striatum.

### 3.2. SCF+G-CSF treatment increases Olig2 positive cells in the ipsilateral external capsule in the chronic phase of severe TBI

Oligodendrocyte transcription factor 2 (Olig2) expressed in both the oligodendrocyte progenitor cells (OPCs) and mature oligodendrocytes plays an important role in the generation of oligodendrocytes [31,32]. Myelin sheath is comprised of oligodendrocyte processes. Myelination is largely dependent on oligodendrocytes [33]. The increases in OPCs and oligodendrocytes are indicators of a cellular response for remyelination after TBI [34,35]. To determine whether oligodendrocyte generation is involved in the SCF+G-CSF-enhanced remyelination in the white matter in the chronic phase of severe TBI, we quantified the number of Olig2^+^ cells in the ipsilateral external capsule over the time course. We observed that the number of Olig2^+^ cells in the TBI-SCF+G-CSF group was significantly increased at 1 day, 21 weeks, and 36 weeks post-treatment compared with the TBI-vehicle group (Figure 4 A and B), suggesting that the SCF+G-CSF-induced persistent increases of Olig2^+^ cells in the ipsilateral external capsule may play a reparative role in enhancing remyelination in the chronic phase of severe TBI.

**Figure 4.**
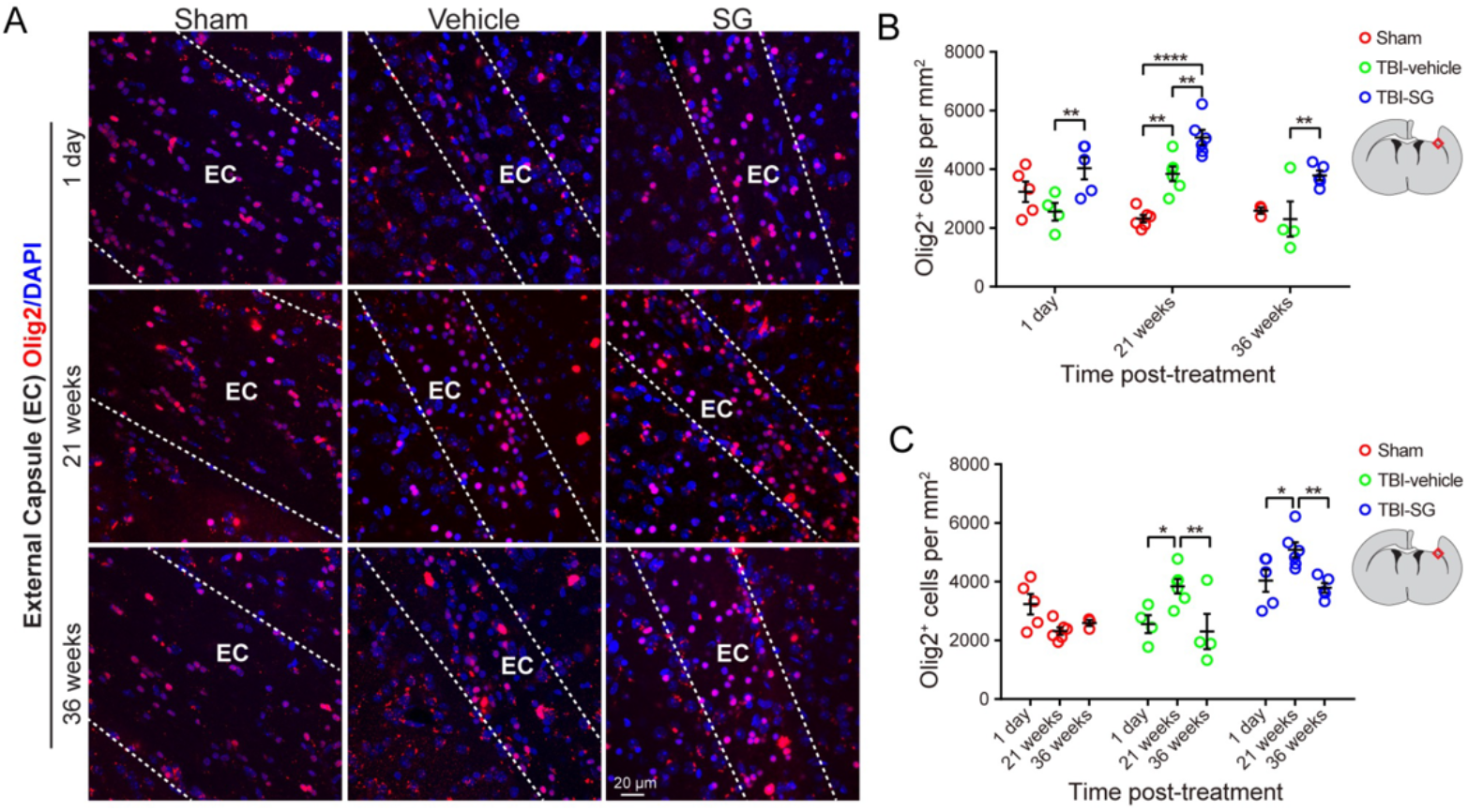
SCF+G-CSF treatment increases Olig2-immunopositive cells in the ipsilateral external capsule in the chronic phase of severe TBI. (**A**) Representative images of Olig2-immunopositive oligodendrocyte progenitor cells and oligodendrocytes in the ipsilateral external capsule at 1 day, 21 weeks, and 36 weeks post-treatment. (**B**) Quantitative data of analysis of the Olig2-immunopositive cells in the ipsilateral external capsule at 1 day, 21 weeks, and 36 weeks post-treatment in the sham, TBI-vehicle, and TBI-SCF+G-CSF groups. (**C**) Quantitative data of analysis of the Olig2-immunopositive cells in the ipsilateral external capsule in the sham, TBI-vehicle, and TBI-SCF+G-CSF groups at different time points. Two-way ANOVA followed by Tukey *post hoc test*. \**p* < 0.05, \*\**p* < 0.01, \*\*\*\**p* < 0.0001. Scale bar: 20 μm. SG: SCF+G-CSF.

We also observed interesting findings showing that at 21 weeks post-treatment, the number of Olig2^+^ cells in both the TBI-vehicle control and TBI-SCF+G-CSF groups was significantly greater than in the sham control group, while the TBI-SCF+G-CSF group showed the greatest increases of Olig2^+^ cells in the ipsilateral external capsule compared with both the sham and TBI-vehicle control groups (Figure 4 A and B), suggesting that the TBI-caused increases of Olig2^+^ cells at this particular time are further enhanced by SCF+G-CSF treatment. We also found that in both the TBI-vehicle and TBI-SCF+G-CSF groups, the number of Olig2^+^ cells at 21 weeks post-treatment was significantly greater than those at 1 day and 36 weeks post-treatment (Figure 4 A and C). These findings suggest that the high number of Olig2^+^ cells appearing at 21 weeks post-treatment (i.e. 34 weeks after TBI) may play a role in maintaining myelin at this time point, and that the decreased number of Olig2^+^ cells showing at 36 weeks post-treatment (i.e. 49 weeks after TBI) could be responsible for the big fall of myelin (severe demyelination) (Figure 2) in this late chronic phase of severe TBI. Our observation indicates that the Olig2^+^ cells in the ipsilateral external capsule are dynamically changing in the chronic phase of severe TBI, which is aligned with the processes of progressive demyelination in the chronic phase of severe TBI.

### 3.3. SCF+G-CSF treatment increases oligodendrocyte progenitor cells in the subventricular zone in the chronic phase of severe TBI

Neural stem cells (NSCs) have been reported to contribute to brain remodeling following TBI [36]. The proliferation of progenitor cells in the subventricular zone (SVZ) is vigorously increased days after TBI, and the progenitor cells migrate from the SVZ to the injury sites in response to TBI [37]. As mentioned earlier, the increase in OPCs is a cellular response to remyelination after TBI. We then sought to determine the efficacy of SCF+G-CSF treatment in increasing OPCs in the SVZ of the TBI brain in the chronic phase. To do so, Olig2 and Sox2 antibodies were used for detecting the OPCs in the SVZ. Sox2 is a marker for both NSCs and OPCs [38,39]. We found that the number of Olig2/Sox2 double-positive OPCs in the contralateral SVZ of SCF+G-CSF-treated TBI mice was significantly greater than in the sham and TBI-vehicle control groups at 1-day post-treatment (Figure 5 A and B). At 21 weeks post-treatment, the number of Olig2^+^/Sox2^+^ OPCs in the ipsilateral SVZ of the SCF+G-CSF-treated TBI mice was significantly increased compared with the mice in the TBI-vehicle control group (Figure 5 A and C). However, there were no significant differences among all groups at 36 weeks post-treatment (Figure 5 A and D). Altogether, our findings suggest that SCF+G-CSF treatment increases the number of OPCs in the SVZ in the chronic phase of severe TBI, and that this effect could maintain for more than 5 months.

**Figure 5.**
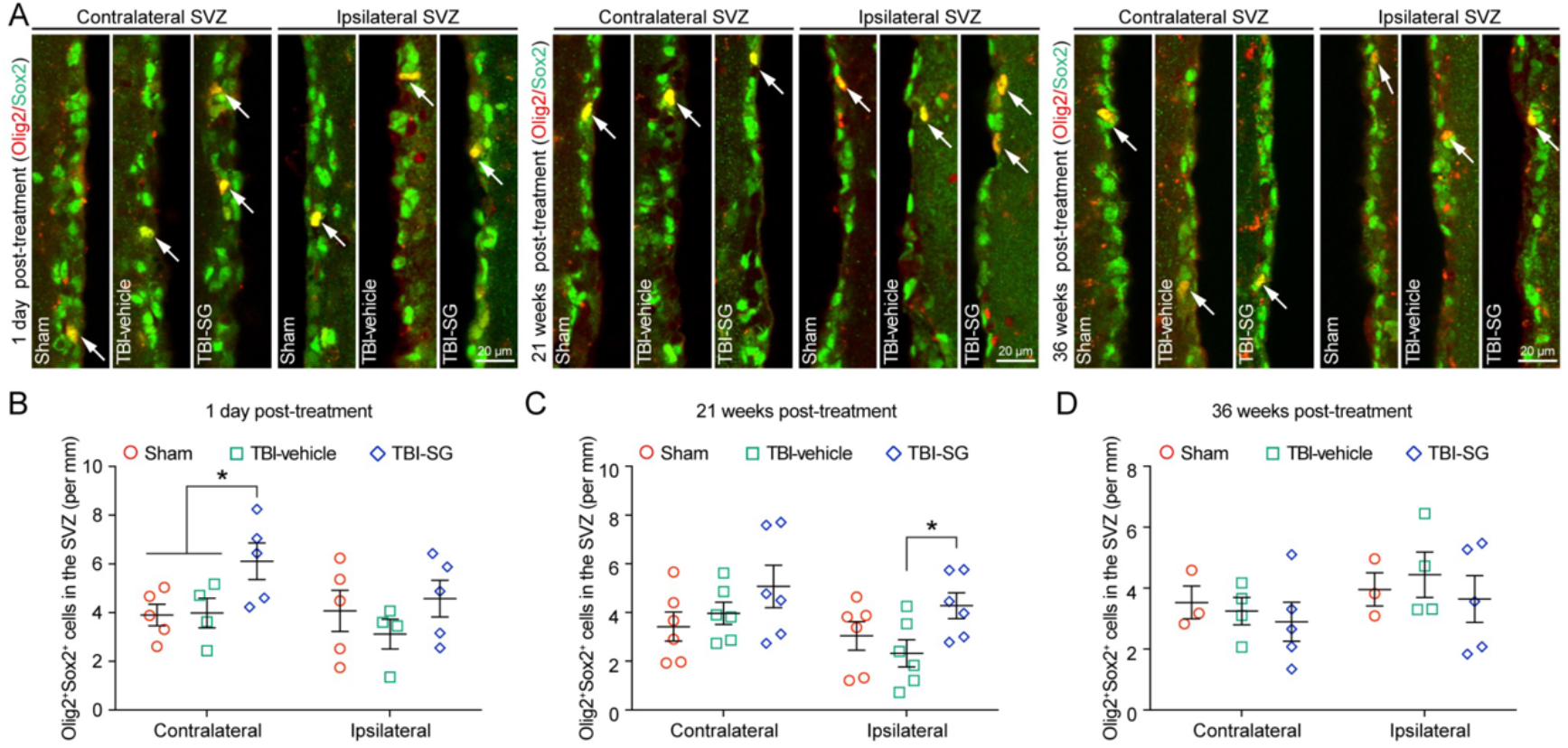
SCF+G-CSF treatment increases the number of oligodendrocyte progenitor cells (OPCs) in the subventricular zone (SVZ) in the chronic phase of severe TBI. (A) Representative images of Olig2^+^Sox2^+^ double-positive OPCs (the arrow-indicated yellow cells) in the contralateral and ipsilateral SVZ at 1 day, 21 weeks, and 36 weeks post-treatment. **(B - D)** Quantitative data of analysis of the number of OPCs in the SVZ at 1 day **(B),** 21 weeks **(C),** and 36 weeks **(D)** post-treatment in three experimental groups. Two-way ANOVAfollowed by Fisher’s LSD test, \**p* < 0.05. Scale bars: 20 μm. SG: SCF+G-CSF.

To validate these findings, we performed double immunofluorescence staining using Ki67 (a proliferation marker) and Olig2 antibodies to detect the proliferation of OPCs in the SVZ at 1-day post-treatment. In both the contralateral and ipsilateral SVZ, the number of Olig2/Ki67 double immunopositive cells was significantly increased in the TBI-SCF+G-CSF group compared with the TBI-vehicle and sham control groups (Figure 6 A-C), suggesting robust effects of SCF+G-CSF treatment in enhancing the proliferation of OPCs in the bilateral SVZ in the chronic phase of severe TBI.

Correlation analyses demonstrated that the number of Olig2^+^/Ki67^+^ proliferating OPCs in the ipsilateral SVZ showed a positive correlation with the percentage of MBP^+^area in the ipsilateral external capsule (Figure 6D) but it did not reach a statistically significant level (p=0.0928). However, there was a significantly positive correlation between the number of Olig2^+^/Ki67^+^ proliferating OPCs in the ipsilateral SVZ and the percentage of MBP^+^ area in the ipsilateral striatum (Figure 6E), indicating that the proliferation of OPCs in the SVZ is linked to the remyelination in the white matter during the chronic phase of severe TBI. SCF+G-CSF-enhanced proliferation of OPCs in the SVZ could contribute to increased remyelination in the white matter of the TBI brain in the chronic phase.

**Figure 6.**
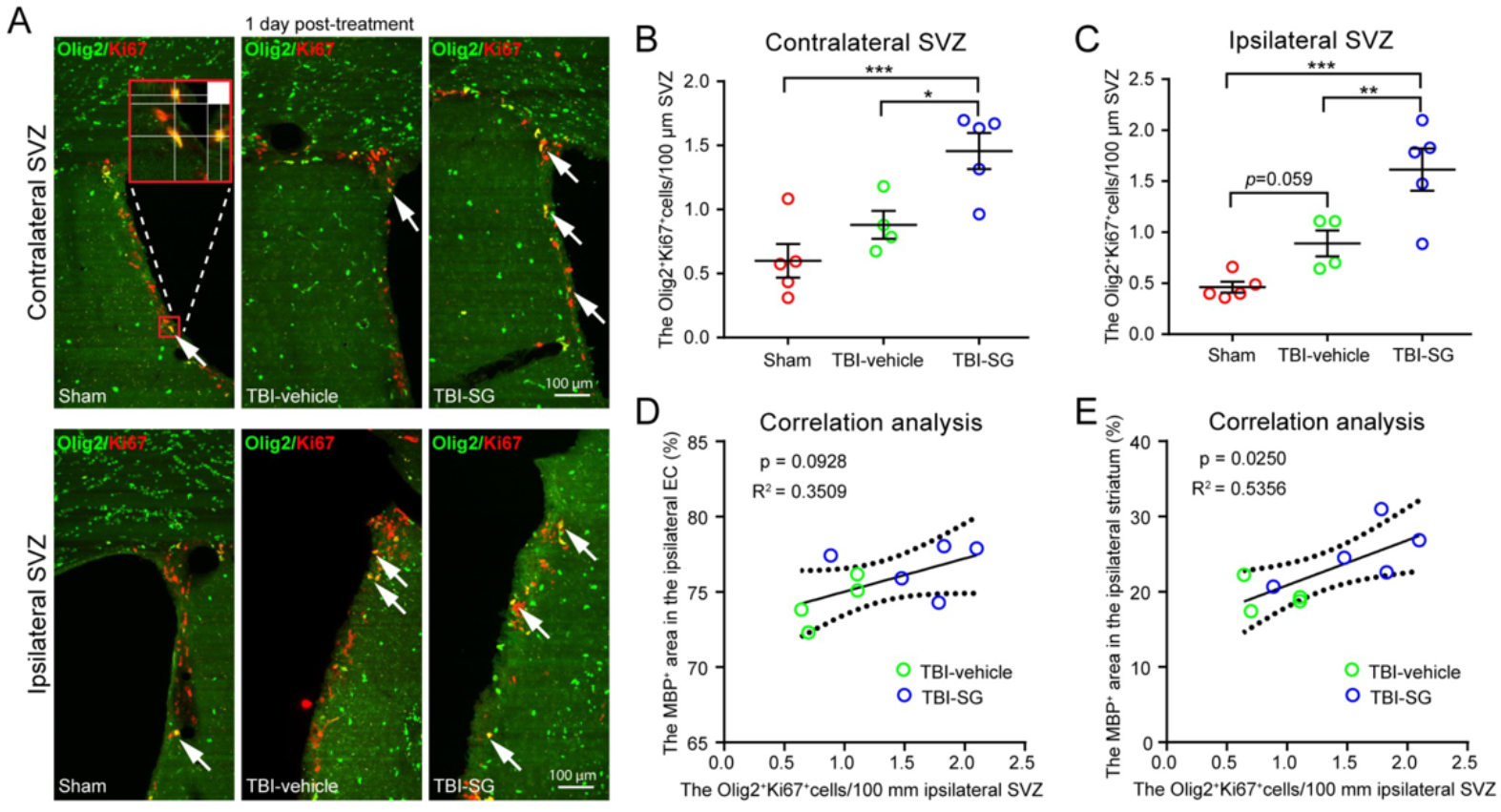
SCF+G-CSF treatment promotes the proliferation of oligodendrocyte progenitor cells (OPCs) in the subventricular zone (SVZ) in the chronic phase of severe TBI. (**A**) Representative images of proliferating OPCs (the arrow-indicated yellow cells coexpressing Olig2 and Ki67) in the contralateral and ipsilateral SVZ at 1 day post-treatment. (B and C) Quantitative data of analysis of the number of Olig2^+^Ki67^+^ double-positive OPCs in the contralateral SVZ **(B)** and Ipsilateral SVZ **(C)** at 1 day post-treatment. One-way ANOVA followed by Fisher’s LSD test, \**p* < 0.05, \*\**p* < 0.01, \*\*\**p* < 0.001. Scale bars: 100 μm. SG: SCF+G-CSF. **(D)** Correlation between MBP^+^ area in the ipsilateral external capsule (EC) and the number of Olig2^+^Ki67^+^ double-positive OPCs in the ipsilateral SVZ. (E) Correlation between MBP^+^ area in the ipsilateral striatum and the number of Olig2^+^Ki67^+^ double-positive OPCs in the ipsilateral SVZ. Pearson’s correlation coefficient test (linear regression) was used for correlation analysis.

### 3.4. SCF+G-CSF treatment promotes oligodendrocyte progenitor cell proliferation and differentiation *in vitro*

To validate the *in vivo* data, we carried out an *in vitro* study through which we sought to examine the efficacy of SCF+G-CSF treatment in promoting the proliferation and differentiation of cultured primary OPCs.

Before starting the experiment, we evaluated the purity of the cultured OPCs using OPC markers. As shown in Figure 7-A and B, the percentages of NG2^+^ cells, Olig2^+^cells, and Sox2^+^ cells were 97.2%, 96.6%, and 94.5% respectively, suggesting that cultured primary OPCs with high purity that can be used for further study. The proliferation of OPCs was assessed 24 h after SCF+G-CSF treatment. The OPCs treated with SCF+G-CSF showed significant increases in the percentage of BrdU^+^/Olig2^+^ cells (Figure 7C and D) and Ki67^+^/Olig2^+^ cells (Figure 7C and E), indicating that SCF+G-CSF treatment can enhance OPC proliferation. We then determined the efficacy of SCF+G-CSF treatment on OPC differentiation. OPCs were cultured in the OPC differentiation medium (i.e. OPC medium without OPC growth factors). As a positive control, some OPCs were treated with T3. After SCF+G-CSF or T3 treatments for 48 h, the differentiated oligodendrocytes were detected by immunofluorescence staining of MBP. The percentage of MBP^+^ cells in T3-treated OPCs was significantly increased compared with medium controls and SCF+G-CSF treatment (Figure 7F and G). Furthermore, SCF+G-CSF treatment significantly increased the percentage of MBP^+^ cells compared to the medium controls (Figure 7F and G), suggesting that SCF+G-CSF treatment can also promote the differentiation of OPCs into oligodendrocytes.

**Figure 7.**
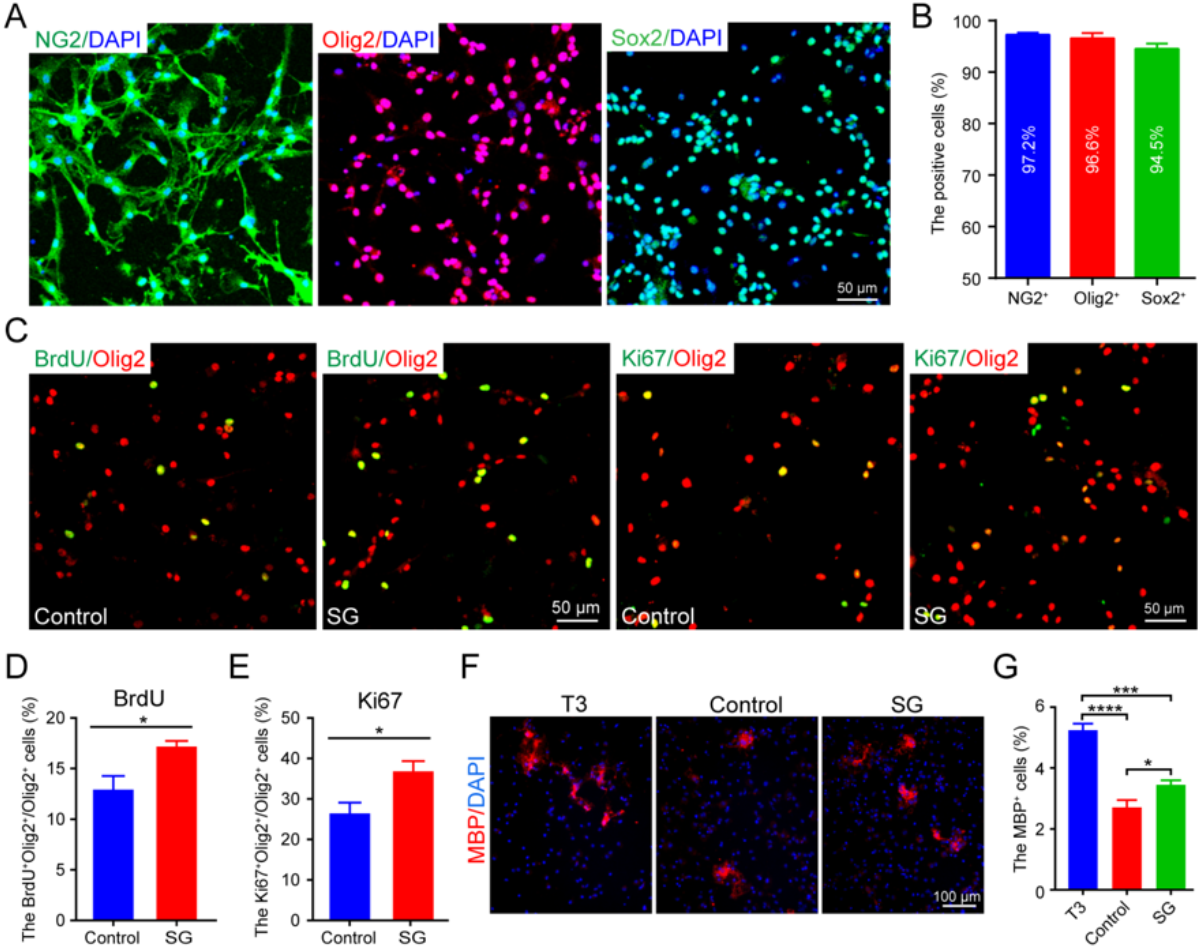
SCF+G-CSF treatment promotes the proliferation and differentiation of oligodendrocyte progenitor cells (OPCs) *in vitro*.**(A)** Representative images show that the primary cultured OPCs express NG2, Olig2, and Sox2. DAPI: *nuclear counterstain*. **(B)** The bar graph shows the percentage of NG2^+^, Olig2^+^ and Sox2^+^ cells in the primary cultured OPCs, indicating the high purity of the cultured OPCs. **I**Sample size: n = 4. (C) Representative images show the proliferating OPCs (BrdU^+^ Olig2^+^ or Ki67^+^ Olig2^+^ double-positive cells) at 24 h posttreatment. Scale bars in panels A and C: 50 μm. (D and E) Quantitative data of analysis of the number of BrdU^+^/Olig2^+^ (D) and Ki67^+^/Olig2^+^ (E) proliferating OPCs in the medium control and SCF+G-CSF treatment groups. Unpaired Student’s t-test, *p < 0.05, n =4. **(F)** Representative images show the differentiated oligodendrocytes (MBP^+^) 48 h after treatment. Scale bar: 100 μm. (G) A bar graph shows the percentage of MBP^+^ oligodendrocytes in T3 treatment (positive control), medium control, and SCF+G-CSF treatment groups 48 h after treatment. SG: SCF+G-CSF. One-way ANOVA followed by Fisher’s LSD test, \**p* < 0.05, \*\*\**p* < 0.001, \*\*\*\**p* < 0.0001, n =4.

## 4. Discussion

Our study has demonstrated that persistent and progressive myelin loss with a temporary increase of oligodendrocytes exists in the chronic phase of severe TBI. The combination of SCF and G-CSF treatment in the chronic phase of severe TBI attenuates the TBI-induced myelin loss and enhances remyelination in the ipsilateral external capsule and striatum, which is associated with SCF+G-CSF-enhanced OPC proliferation and oligodendrocyte generation. Myelin impairments disrupt the axonal integrity and neuronal signaling conduction, which may lead to cognitive decline, motor dysfunction, and emotional symptoms commonly observed after TBI [40,41]. It is plausible that SCF+G-CSF-improved neurological function recovery in chronic TBI [23][27] could be linked to the enhancement of remyelination in the chronic phase of severe TBI. Our findings reveal the potential role of OPCs and oligodendrocytes in SCF+G-CSF-enhanced myelin repair in the chronic phase of severe TBI.

White matter contains myelinated axons which are highly vulnerable to damage after TBI. White matter injury is associated with long-term impairments of neurological function after TBI [42]. Myelin pathology is the key feature of white matter injury post-TBI. Post-TBI demyelination that has been observed in both humans [43] and rodent TBI models [13] is linked to long-term deficits in sensorimotor and cognitive function [40,16]. Since most experimental TBI studies target demyelination/remyelination in the white matter days and weeks after TBI [34,16,13,44,45], it remains largely unknown whether white matter demyelination occurs months after severe TBI in the chronic phase. Our findings from examining the MBP expression in the ipsilateral external capsule and striatum indicate that progressive demyelination in the white matter persists up to more than one year after severe TBI, which exhibits an exacerbating myelin loss at 12.25 months post-severe TBI. It has been shown that the apoptosis of oligodendrocytes starts in the acute phase of TBI and remains at least three months post-TBI [14,13][46], which may contribute to the persistent and progressive demyelination in the chronic phase of TBI. It has been thought that chronic demyelination is an irreversible pathology leading to progressive functional decline [34]. In the present study, our data have revealed that treatment with SCF+G-CSF in the chronic phase of severe TBI could reverse the chronic demyelination by enhancing remyelination, which provides the feasibility of repairing the damaged myelin in the chronic phase of TBI and reveals a long-term therapeutic window for myelin repair after TBI.

The myelin sheath is comprised of oligodendrocyte processes. Myelination is largely dependent on oligodendrocytes [33]. Increased OPCs and oligodendrocytes are the indicators of a cellular response for remyelination after TBI [34,35]. Olig2 is the oligodendrocyte lineage transcription factor and is expressed in both the OPCs and oligodendrocytes [47,48]. In the present study, we found that the number of Olig2^+^ cells in the ipsilateral external capsule was increased 1 day, 21, and 36 weeks after SCF+G-CSF treatment in the chronic phase of severe TBI. This observation indicates a longterm effect of SCF+G-CSF treatment in enhancing oligodendrocyte generation in the white matter tract, which lays a vital foundation to support remyelination in the chronic phase of severe TBI. Our findings also showed that the number of Olig2^+^ cells in the ipsilateral external capsule was increased at 34 weeks post-TBI (i.e. 21 weeks posttreatment) and then dropped down at 12.25 months post-TBI (i.e. 36 weeks posttreatment) in all TBI mice. This dropping down could be attributed to reactive oligodendrogenesis exhausting during the long-term stimulation of demyelination [49–51] or the interaction with age-related decline in oligodendrogenesis [52]. Interestingly, MBP^+^ myelin also dramatically dropped down in the ipsilateral external capsule at 12.25 months post-TBI for all TBI mice compared to the level shown at 34 weeks post-TBI, which is consistent with the dropping down pattern of Olig2^+^ cells in the same region. These findings confirm the tight link between the ability of oligodendrogenesis and the severity of post-TBI demyelination.

The SVZ is a persistent source of OPCs in the adult brain. The OPCs in the SVZ of the adult brain can proliferate, migrate, and differentiate into mature oligodendrocytes which restore impaired myelin sheaths in injuries and diseases of the central nervous system (CNS) [53–55]. Studies in both human severe TBI and animal models of TBI have shown increases in OPCs days, weeks and even up to 3 months after TBI, and the TBI-increased OPCs have been proposed to play a role in myelin repair [13,14,46]. In the present study, we have demonstrated that SCF+G-CSF treatment initiated at 3 months after severe TBI promotes OPC proliferation and increases the number of OPCs in the SVZ at 1 day post-treatment. It has been reported that SCF+G-CSF injections during 11-20 days after spinal cord injury in mice promote the proliferation of OPCs [26]. OPCs express the receptor of SCF named C-KIT which loses its expression in OPCs when OPCs differentiate into mature oligodendrocytes [56], while the G-CSF receptor is expressed in the oligodendrocytes [25], indicating that SCF and G-CSF may play different roles in OPC proliferation and differentiation for myelin repair. The data from our *in vitro* study further confirmed the direct role of SCF+G-CSF in promoting OPC proliferation and differentiation. SCF and G-CSF may also be indirectly involved in myelin repair by improving the microenvironment, such as anti-inflammation, neuroprotection, and brain-derived neurotrophic factor (BDNF) production [25,17,57,23,58]. It has been shown that G-CSF protects oligodendrocytes from death via the suppression of inflammatory cytokines and up-regulation of anti-apoptotic proteins in spinal cord injury [25]. Taken together, SCF and G-CSF may enhance myelin repair after CNS injury through direct and indirect mechanisms.

Our previous studies have demonstrated that SCF+G-CSF treatment in both the subacute phase and the chronic phase of TBI improves neurological function recovery and ameliorates pathological progression post-injury [27,23]. The present study provides additional evidence supporting the long-term effects of the SCF+G-CSF treatment on myelin repair after severe TBI. However, the molecular mechanism underlying the SCF+G-CSF-enhanced myelin repair in the chronic phase of severe TBI remains unclear. Our previous studies have shown that SCF+G-CSF treatment activates MEK/ERK/p53 and PI3K/AKT/NF-κB/ BDNF signal pathways to enhance neurite growth [24,57]. Further studies need to be performed to elucidate whether the same molecular mechanism is involved in the SCF+G-CSF-enhanced myelin repair, and whether SCF+G-CSF treatment could be applied in other demyelination-related diseases.

## 5. Conclusions

In the present study, we demonstrate that persistent myelin impairment exists in the chronic phase of severe TBI and that SCF+G-CSF treatment in the chronic phase of severe TBI enhances remyelination in the ipsilateral white matter. Furthermore, SCF+G-CSF treatment in the chronic phase of severe TBI promotes the proliferation of OPCs in the SVZ and increases OPCs and mature oligodendrocytes in the ipsilateral white matter, suggesting that the OPCs in the SVZ may migrate to the demyelinated sites and differentiate into mature oligodendrocytes to repair the TBI-induced demyelination.

## Acknowledgements

We are grateful to Karen Hughes for her assistance in proofreading the manuscript.

